# Assessing the use of RIVPACS-derived invertebrate taxonomic predictions for river management

**DOI:** 10.1101/2024.06.14.599001

**Authors:** A. Johnson, J. Murray-Bligh, L.E. Brown, A.M. Milner, M.J. Klaar

## Abstract

The River Invertebrate Prediction and Classification System (RIVPACS) is used widely in freshwater management to set targets for macroinvertebrate ecological health based on the expected scores of metrics such as WHPT or LIFE in the absence of anthropogenic stressors. An underutilised capability of RIVPACS-type models is the capability to predict expected macroinvertebrate community composition, which could function as a novel management metric for river health. We present a novel Monte-Carlo simulation approach that generates simulated expected communities for England’s rivers based on RIVPACS predictions. This allows for assessments of macroinvertebrate health using similarity calculations between observed and expected communities. We assess 10-year trends in similarity between 2010 and 2019 at 4172 sites in England, and contrast these trends with WHPT ASPT O/E trends in the same period. Similarity scores include both Chi-Squared and Hellinger methods, to prioritise rare and common species, respectively. We find that whilst most sites (63.3%) showed improvement in WHPT ASPT O/E in this period, most sites showed declines in similarity for Chi-Squared and Hellinger O/E (51.1% and 58.8%, respectively). We identified three case study regions showing contrasting trends and illustrate how the new RIVPACS-derived similarity calculations can track meaningful shifts in composition associated with water quality and multiple stressors including invasive species. RIVPACS-derived similarity calculations potentially provide a sensitive and practical management metric to assess ecosystem health, although further work is required to understand the composition of communities in changing environments with clear changes in stressor regimes.

## Introduction

The River Invertebrate Prediction and Classification System (RIVPACS) is a tool used in freshwater management, allowing practitioners to assess the ecological health of rivers by comparing observed metric scores to expected targets in the absence of anthropogenic pressures (Clarke *et al*., 2003; Wright *et al*., 1989). Originally developed in the UK, variations of the RIVPACS approach are now used globally, and have been incorporated in the European Union’s Water Framework Directive as a method for setting ecological quality targets and allowing comparison of degradation between sites (European Union, 2000; Davy-Bowker *et al*., 2006). RIVPACS-type models have been developed in Canada (CABIN), Australia (AUSRIVAS), and Spain (MEDPACS) (Manual, 2009; Smith *et al*., 1999; Poquet *et al*., 2009). In the UK, RIVPACS is used primarily to establish targets for metrics such as the Walley, Hawkes, Paisley & Trigg (WHPT) and Lotic Invertebrate Flow Evaluation (LIFE) scores (Chen *et al*., 2019), tracking pollution and general river degradation, and flow modification effects, respectively. However, these metrics provide only narrow and incomplete assessments of ecosystem health and are not capable of providing insights into underlying community dynamics. Additionally, some metrics can co-vary, and thus it can be difficult to distinguish between underlying stressors (Jones et al., 2023). These metrics have also been shown as insensitive to larger trends such as the impact of climate change, or invasive species (Mathers *et al*., 2016; Thompson *et al*., 2018). However, an underutilised capability of RIVPACS-type models is the option to predict macroinvertebrate assemblage composition in the absence of anthropogenic stressors. Using taxonomic similarity as a metric of ecosystem health might allow novel techniques to detect underlying changes that traditional metrics are not designed to detect, such as invasive species or ramp pressures such as climate change. However, it is first necessary to understand whether taxonomic similarity calculations show consistent change in improving or worsening environments, to determine whether such metrics are feasible.

RIVPACS-type models were developed originally to predict macroinvertebrate assemblages that might be found in sites with minimal anthropogenic pressures (Wright, 2000). Communities at minimally impacted reference sites are clustered into unique groupings and compared to new sites based on habitat characteristics (Reynoldson and Wright, 2000; Reynoldson *et al*., 2014). These clusters were generated from 421 reference sites in England using a TWINSPAN clustering algorithm, paired with a linear discriminant analysis to distinguish between clusters based on habitat characteristics. The reference dataset contains approximately 2000 macroinvertebrate samples collected between 1978 and 2002. Since the initial implementation, RIVPACS has been adapted to produce metric targets for sites, based on weighted averages derived from the probability of existing in each cluster rather than using the taxonomic predictions directly, to save computational power by reducing the need to calculate entire simulated communities (Paisley *et al*., 2007; 2014). These metrics are associated with various habitat associations and stressor tolerances of macroinvertebrate taxa, often derived either from expert opinion or statistical studies. Whilst accessible and simple, this approach has many limitations. The individual metrics are often linked to functional traits, such as body size, feeding group, and respiratory capability but there is significant autocorrelation among taxon metric scores (McKenzie *et al*., 2022). Similarly, functional redundancy between taxa means that these metrics might be insensitive to underlying ecosystem dynamics, such as shifts in extinction or invasion rates, or natural turnover (Gillespie *et al*., 2020). Taxonomic targets derived from pre-existing RIVPACS-type models could allow novel assessments of ecosystem health, including taxonomic similarity, as well as allowing the establishment of targets for ecosystem restoration. Similarly, taxonomic targets can be expected to be sensitive to pressures currently unaccounted for in traditional metrics, such as invasive non-native species or extreme climate events.

RIVPACS is often referred to as a classification-first community predictive model (CPMs). This approach differs from Species Distribution Models (SDMs), which predict the habitat preferences of individual species or taxa and estimate their abundance based on site conditions (Miller, 2010). In contrast, CPMs predict the abundances of entire communities simultaneously. Instead of modelling the abundance or likelihood of specific taxa, RIVPACS forecasts the likelihood of known average community compositions. It then assigns likelihoods to individual taxa based on the average abundances observed within each compositional cluster. At new sites, the recorded environmental conditions are used to generate predicted abundance likelihood distributions for each taxon, facilitating the prediction of expected community compositions. Two European studies developed custom RIVPACS-type models for macroinvertebrates and lake phytoplankton in Spain and Norway, respectively, finding that the models performed well for predicting the presence/absence of common taxa, but struggled at predicting rarer taxa (Feio *et al*., 2014; Hallstan *et al*., 2012). In contrast, an assessment of RIVPACS-type models and species models for fish assemblages in Australia found that the RIVPACS-type model had lower predictive capabilities (Rose *et al*., 2016). A study in the USA furthermore demonstrated the potential of using Bray-Curtis distances between an average expected community and observed communities, but did not use trends over time or large-scale simulations to account for variation in the range of possible communities at a site (Van Sickle, 2008). To date there has been no assessment of RIVPACS taxonomic capabilities for analysing long-term trends or large-scale spatial dynamics of invertebrate biodiversity. Such an analysis is vital to determine whether RIVPACS-type models are suitable for setting taxonomic targets for macroinvertebrate communities. To assess the feasibility of RIVPACS-type taxonomic targets as a management metric, it is necessary to determine:

1. if target compositions are achievable in a changing environment;
2. how the composition of sites that are experiencing changes in multiple-stressor changes relative to their target compositions, and;
3. whether or not taxonomic trends are sensitive to novel stressors such as climate change or invasive species.

This work evaluates the potential use of RIVPACS derived taxonomic targets for use as river ecosystem management metrics, investigating whether underlying trends can be observed, and how this provides alternative information to traditionally used river biodiversity health metrics. To accomplish this, a novel Monte Carlo simulation was produced which is capable of integrating into the RIVPACS model to generate simulated community assemblages. This simulation was then used to assess 10-year trends in taxonomic similarity across England for the 2010-2019 period. These trends are contrasted with trends in the commonly used WHPT ASPT metric, and spatial patterns mapped to assess the sensitivity of taxonomic trends to both multi-stressor impacts and novel stressors.

## Methodology

To date, studies seeking to unpack the macroinvertebrate predictive capabilities of RIVPACS have been limited by the computational power required to accurately model the range of possible compositions allowed by the model (Logez *et al*., 2019). Rather than providing a single predicted composition, RIVPACS produces a range of log-likelihoods for all UK freshwater macroinvertebrate taxa in each log-abundance category. Each community has specific probabilities of arising, based on the abiotic conditions of the target site. Within this likelihood range can be communities that would not exist due to competition between taxa or might not develop if the necessary taxa are not able to re-establish in the ecosystem. This presents challenges when generating taxonomic targets. If too few target compositions are produced, it is likely this would erroneously increase the distance between target and observed compositions, by comparing real compositions to compositions that are unlikely to exist.

To overcome this, it was necessary to simulate thousands of communities for each individual site, and generate distance calculations for these against a given sample. We developed an expansion to the RIVPACS model that can automatically generate simulated compositions and undertake the necessary distance calculations. Analyses followed three steps:

1. Building a Monte-Carlo simulation (Raychaudhuri, 2008) to allow for similarity calculations between an observed community (sample) and the range of theoretical possible target compositions.
2. Calculating changes over time, as well as spatial patterns, for both invertebrate composition similarity and WHPT ASPT Observed/Expected (O/E) ratios.
3. Investigating spatial autocorrelations to determine potential sensitivity to multi-stressor impacts and novel stressors, including invasive species.

Data processing and management, and non-spatial statistics and simulations were undertaken in R 4.0.2 using RStudio 1.3 (R Core Team, 2018). Spatial statistics were undertaken using GeoDa 1.14 (Anselin *et al*., 2006). The Environment Agency freshwater river macroinvertebrate surveys (Biosys) dataset for English rivers since 2010 served as the primary data source (Environment Agency, 2022). Surveys are carried out with comparable sampling methods at consistent sample sites, allowing trends and shifts to be detected (Murray-Bligh *et al*., 1997). Prior to 2010, data management protocols shifted over time with a change from log10-to actual abundances being recorded, and changes in taxonomic level of identification, preventing longer-term analyses, with a full description of changes provided with the dataset (Environment Agency, 2022).

### Monte-Carlo simulation and WHPT ASPT predictions

To allow for the calculation of trends in WHPT ASPT and taxonomic trends at each individual site, and to ensure comparability between sites, the following selection criteria were used for the trend analysis:

1. Only samples between 2010-2019 were considered to provide a consistent sampling period between sites.
2. Within this year range, a site must have been studied for a duration of at least 5 years, and a minimum of 5 samples collected over the study period to ensure a temporal range for trend analysis.
3. Sites must have a record of habitat characteristics, such as slope and substrate to enable predictions to be generated in the RIVPACS model (Wright, 2000). An exception was for alkalinity and conductivity which can be estimated within the RIVPACS model.

A total of 4172 sites were retained, comprising 33,586 individual samples, and taxonomic records at each site were aggregated to at least family level (Figure 1). Records at higher levels that were not identified in the RIVPACS taxonomic record were excluded. Taxonomic data were then transposed onto the relevant sample data using the consistent sample identification code. WHPT Observed/Expected (O/E) ratio was calculated using the Average Score Per Taxa (ASPT), which accounts for the number of taxa observed at each site.

**Figure 1:**
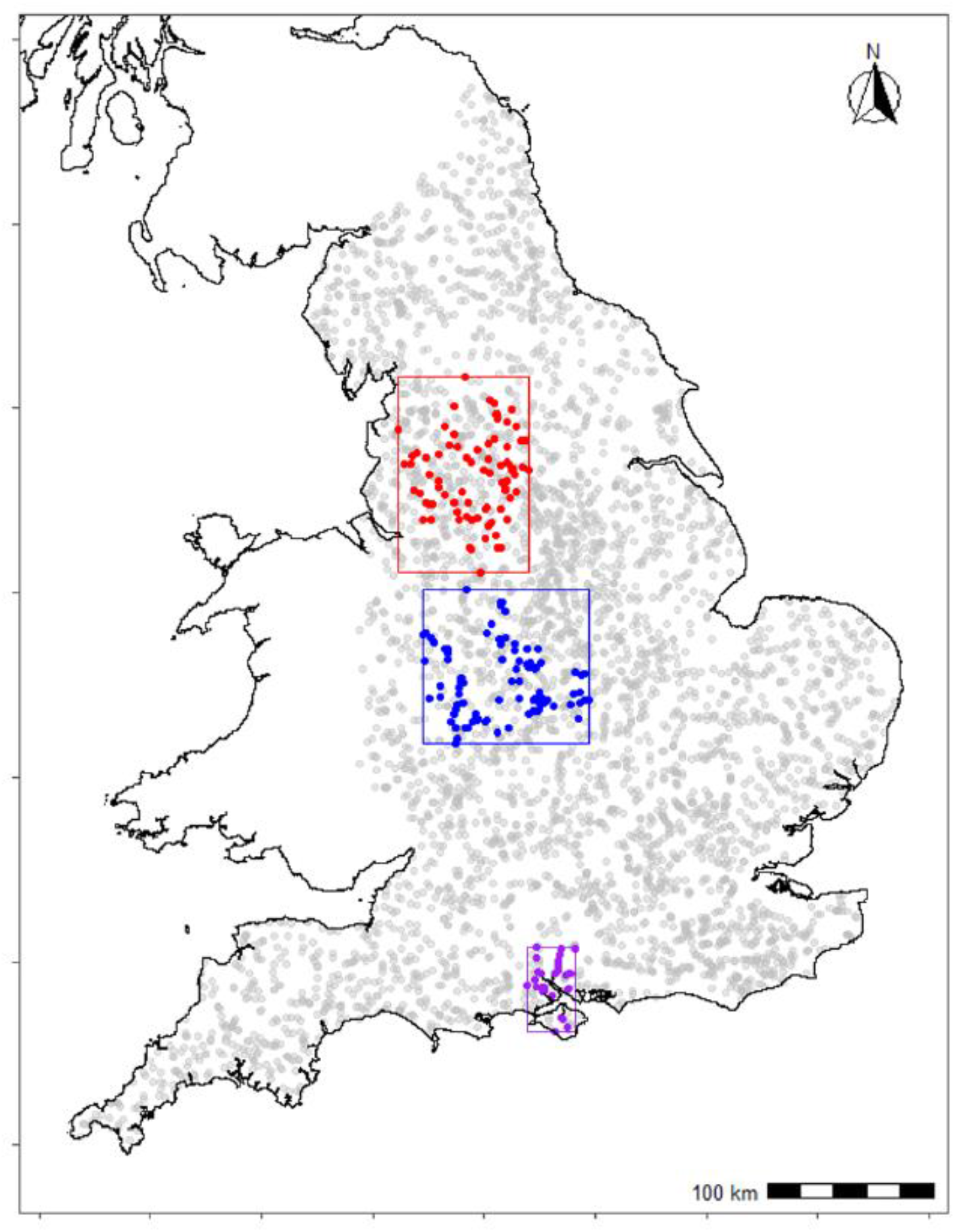
Map of all sites that were included in the trends analysis (grey circles), and the regions selected for case studies (boxes). The Lancashire region (red) showed agreement between WHPT ASPT O/E and both taxonomic trends, the West Midlands region (blue) showed improvements in WHPT ASPT O/E and no change/worsening of taxonomic trends, and the Hampshire (purple), showed worsening in WHPT ASPT O/E and Chi-Squared similarity, but improvements in Hellinger similarity.

RIVPACS taxonomic predictions produce a range of log-abundance category likelihoods for different taxa and for each season at each site, derived from the similarity of habitat characteristics of the site to the predicted clusters. RIVPACS taxonomic predictions were produced at the family level (TL3 in RICT) using BIOSYS site data information as input. RIVPACS codes and support files were obtained through the RICT tool, and modified in R to allow a bulk run of all sites on local directories. All relevant similarity calculations between sample and reference sites and log-abundance likelihoods for each taxon within RIVPACS were left unmodified.

A Monte-Carlo simulation was built to generate predicted invertebrate assemblages for each site. For each sample, the range of log-abundance likelihoods for the appropriate season and site were selected and converted to cumulative probability (Table 1). For each taxon, a random number between 0 and 1 was generated. This was compared with the cumulative probability to assign a log-abundance category (Table 1). This was undertaken 1000 times for each of the represented taxa, producing a matrix of 1000 simulated communities. As is protocol in the Biosys dataset, log-abundance categories were recorded using an integer (i.e. 3 for 1-9, 33 for 10-99). Each sample was paired with the respective simulated community matrix. To facilitate distance calculations, Hellinger and Chi-Squared distance transformation were used as standardisation techniques. Hellinger transformation reduces the influence of rare species and prioritises common species. It is defined as:

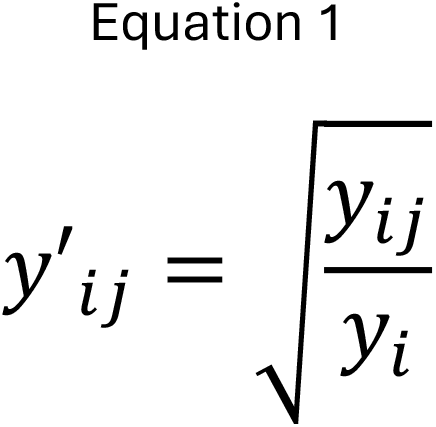

**Table 1.** Summary of the cumulative probability used to convert likelihoods for use in the Monte-Carlo simulation

| Log abundance | Likelihood | Cumulative probability |
| --- | --- | --- |
| 0 | 0.34 | 0.34 |
| 1-9 | 0.19 | 0.53 |
| 10-99 | 0.24 | 0.77 |
| 100-999 | 0.13 | 0.9 |
| 1000-9999 | 0.08 | 0.98 |
| 10000-99999 | 0.02 | 1 |

Where j indexes the species, i the site/sample, and i. is the row sum for the i^th^ sample. Chi-Squared distance transformation increases the influence of rare species and reduces common species. It is defined as:

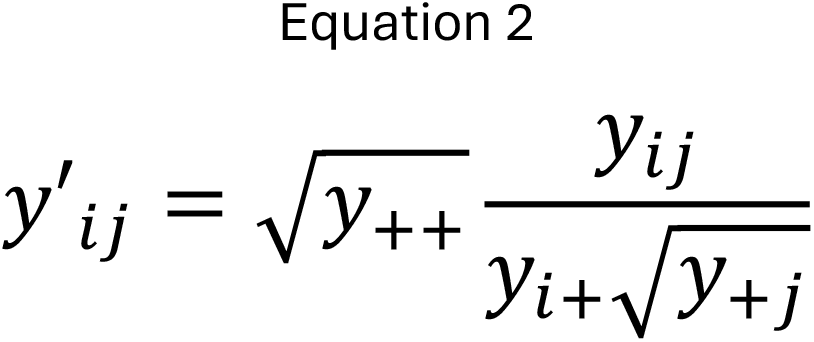

Where y^ij^ is a species presence or abundance value, y^i+^ is the sum of values over row (object) i, y^+j^ is the sum of values over column (species) j, and y++ is the sum of values over the combined community matrix. Both transformations were carried out on independent copies of the community matrix. Pair-wise Bray-Curtis similarity was calculated between the recorded sample and each simulated community. The minimum distance, mean distance, and standard deviation of distances between an observed sample and all simulated samples was recorded for both Hellinger transformed communities and Chi-Squared distance transformed.

Predicted WHPT ASPT values for each site were calculated using RIVPACS code modified within R Studio. This operates in a similar manner to the invertebrate community classification code, calculating likelihoods based off habitat characteristics to all clusters and calculating a likely WHPT ASPT score and confidence. When a target WHPT ASPT prediction was obtained for each site and season, a WHPT ASPT Observed/Expected (WHPT ASPT O/E) was calculated for each individual sample.

### Change over time and spatial statistics

Change over time of both WHPT ASPT O/E and taxonomic similarity were modelled using linear models. For each site, a linear model was calculated for each of the dependent variables with year as independent variable. Slope coefficient was used to infer change in the dependent variable per year.

Spatial trends in rates of change of WHPT ASPT O/E and invertebrate distances were examined independently using a Univariate Moran’s I (UMI) test. Spatial weights were generated using a Euclidian distance with an inverse distance weight of 2. This greatly prioritises closer neighbours over further. A UMI tests correlation of a variable (x) with the mean value of the variable x (⍰) at the nearest neighbours, with means adjusted by the spatial weights. This highlights regions with high levels of spatial autocorrelation. This was then tested against a null hypothesis of no spatial autocorrelation, and significant clusters (p< 0.05) were extracted.

To determine consistent patterns between WHPT ASPT O/E and invertebrate distances, Principal Component Analysis (PCA) was used to pair WHPT ASPT O/E with the respective invertebrate distances. For both PCAs, data were centred and standardised using a z-score transformation. In both instances, the first principal component (PC) represented agreement between trends in WHPT ASPT O/E and invertebrate distances, and the second principal component represented disagreement between trends. To detect spatial trends, UMI’s were used on PC1 and 2 respectively. Clusters were identified for p < 0.05.

Regions that exhibited strong trends in spatial autocorrelation were identified and extracted to investigate underlying drivers and patterns in invertebrate communities. Three regions showing specific patterns in trends were selected, one showing strong widespread agreement in WHPT ASPT O/E and taxonomic similarity, one showing disagreement between WHPT ASPT O/E and both taxonomic similarity metrics, and one showing disagreement between the taxonomic metrics. In each region, the site selection criteria were expanded to include samples from 2000 to 2020, with the first sample at a site having to be collected before 2005, and the last after 2015. This was undertaken to better understand the underlying trends and drivers. To characterise shifts in composition, a Principal Component Analysis was used on the Hellinger transformed invertebrate composition data for all samples at sites within three regions showing strong autocorrelations among WHPT ASPT O/E and invertebrate community composition distances, using the vegan package in R (Oksanen *et al*., 2019). Hellinger transformed data were used to reduce the impact of rarer species that can result in ordination errors. In each region, changes over time were highlighted using ordination ellipses. Samples were grouped into 5-year aggregates from 2000 to 2020. Five-year aggregates were selected as a compromise to highlight key underlying trends whilst ensuring an equal number of sites in each grouping. Shorter period groupings were found to have uneven numbers of samples, which made visualisation of underlying trends more susceptible to sampling effort. An ordination ellipse was calculated for each grouping using 1 standard deviation (SD) in composition expression of the first and second PCA axis. This allowed for a visual representation in the shift of compositions on the main axis. The largest eigenvalues on the first and second axis were identified to detect the key invertebrate taxa that had changed.

## Results

Linear trend analysis for 2010-2019 found that 63.3% of sites have shown an improvement in WHPT ASPT O/E, increasing on average by 0.0032 ± 0.013SD per year. Conversely, only 41.2% of all sites have shown improvement in common-species weighted Hellinger similarity over time. Similarity has changed on average by -0.0023 ± 0.011SD per year, indicating widespread trends away from predicted assemblages overall. In total 49.9% of sites have shown improvement in rare-species weighted Chi-squared similarity over time on average by -0.0001 ± 0.0031SD per year. The greatest decreases in similarity were observed in the south of England and Cornwall (Figure 2). Hellinger and Chi-squared similarity show similar regional patterns with small but notable differences. For example, Norfolk was found to have greater improvements in Chi-Squared similarity compared to Hellinger, and regions in Northumberland with worsening trends in Hellinger similarity performed better in Chi-Squared similarity.

**Figure 2:**
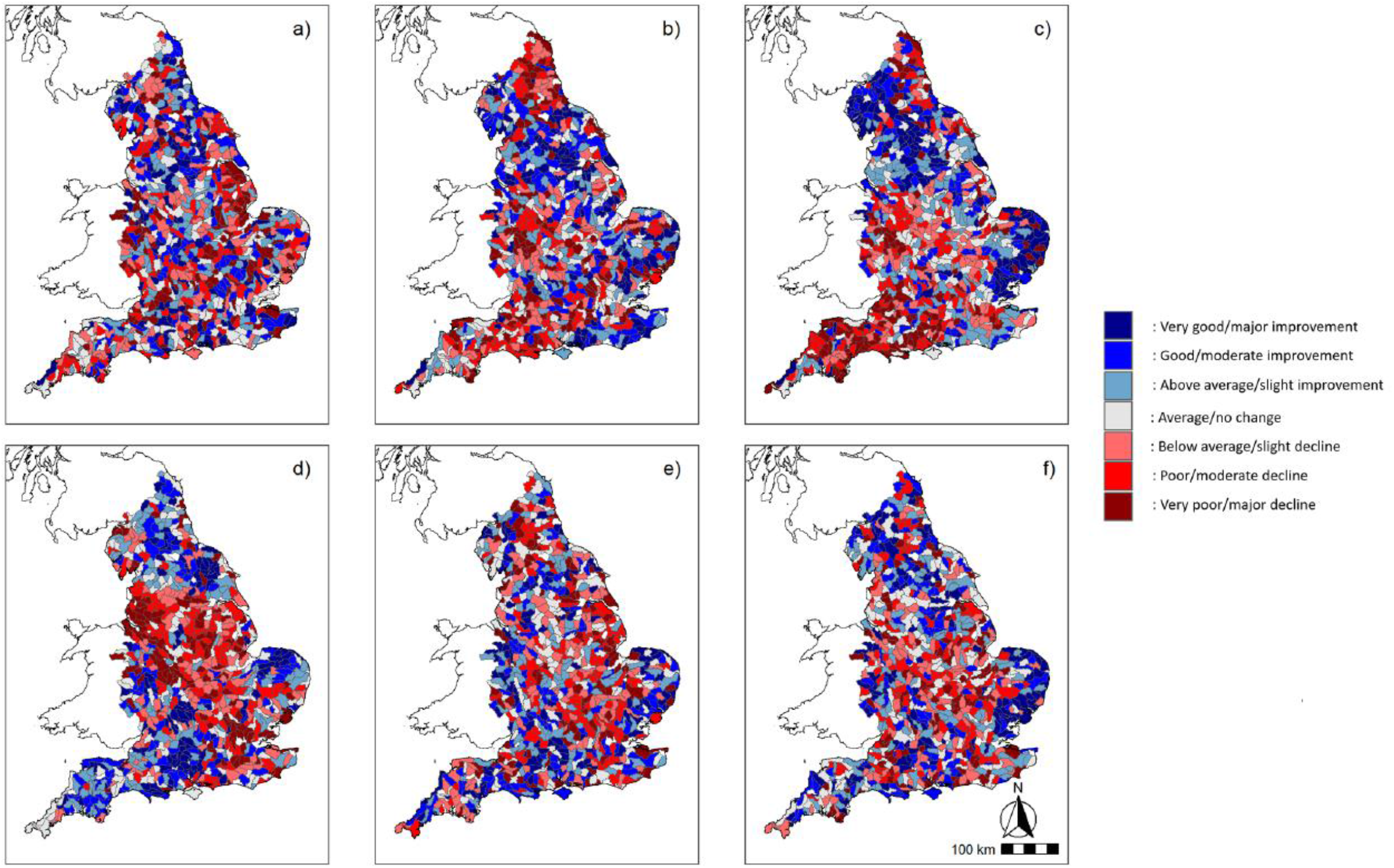
Average change per year in WHPT ASPT O/E (a), Hellinger similarity (b), and Chi-squared similarity (c), and final recorded scores in WHPT ASPT O/E (d), Hellinger similarity (e), and Chi-squared similarity (f) averaged between sites within sub-basins. In all cases, red represents worsening or bad conditions, and blue represents improving or good conditions.

Moran’s I test revealed complex patterns of autocorrelation across England (Figure 3). Manchester and the Peak District have shown consistent patterns of improvement for WHPT ASPT O/E, Hellinger, and Chi-squared similarity. There was partial agreement between Hellinger and Chi-squared similarity in the West Midlands and Shropshire regions, with both showing trends away from predicted assemblages. This was more widespread for Hellinger similarity. Regions of spatial autocorrelation for WHPT ASPT O/E tended to be smaller than the similarity metrics, potentially indicating more spatially discrete management practices or pollution events. A large portion of the south coast of Devon showed consistent decreases in Chi-squared similarity, indicating trends away from RIVPACS predicted communities in this region. This trend was not present for either WHPT ASPT O/E or Hellinger similarity. Similarly, regions in Suffolk and Essex have shown increases in Chi-squared similarity that are not present in WHPT ASPT O/E or Hellinger similarity.

**Figure 3:**
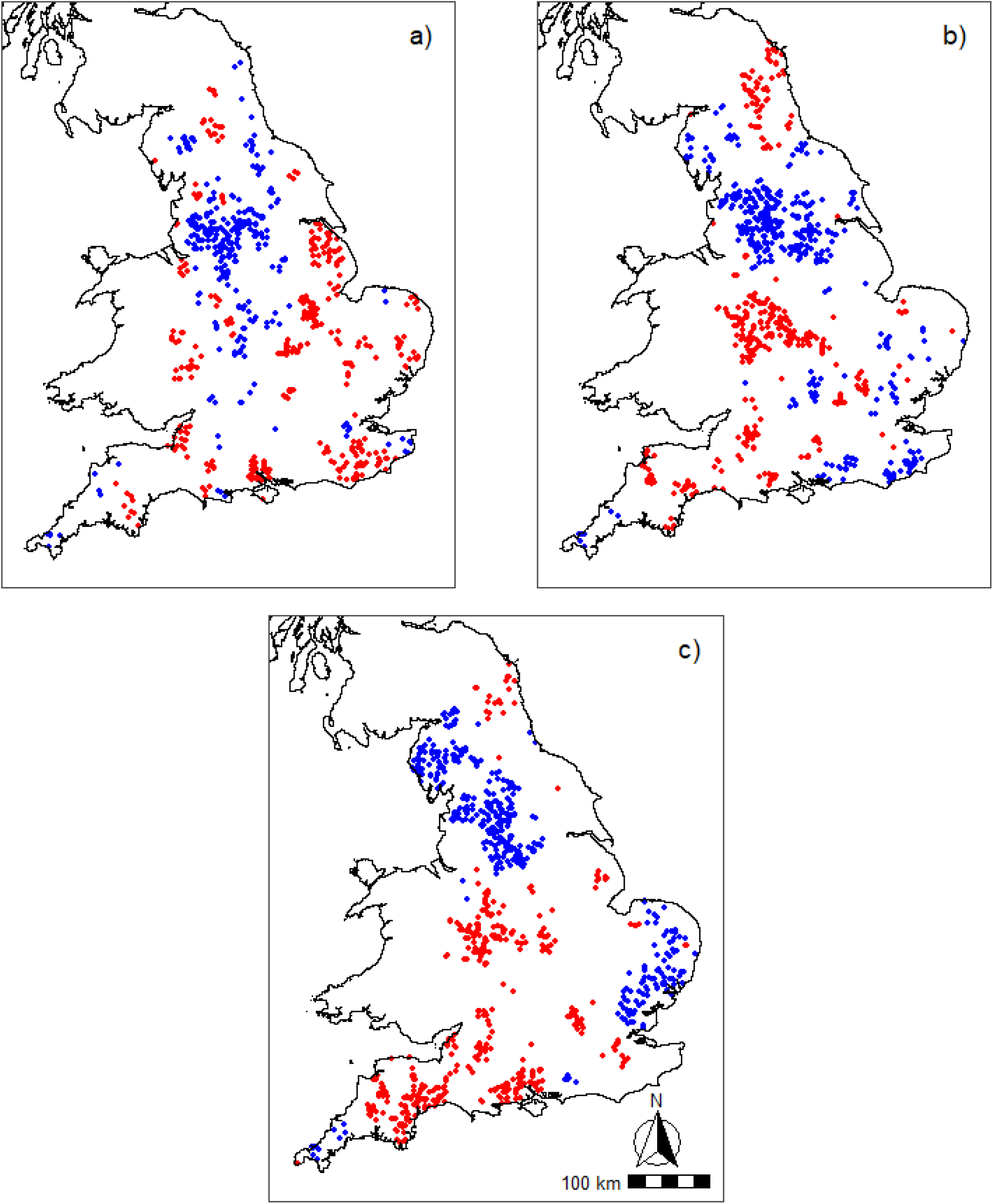
Spatial autocorrelation detected in a) WHPT ASPT O/E, b) Hellinger transformed similarity, and c) Chi-Squared transformed similarity. Blue represents regions showing consistent improvement in the tested metric, and red represents consistent worsening over time

In the PCA of trends in Hellinger and WHPT ASPT O/E, the first PC accounted for 62.3% of the overall variance, and the second accounted for the remaining 37.7%. For Chi-squared and WHPT ASPT O/E, the first principal component accounted for 57.8% of the overall variance. In both circumstances, the first principal component signified regions where the similarity metric agreed with WHPT ASPT O/E (both were improving or worsening), and the second principal component indicated regions with disagreement. Fewer regions of spatial autocorrelation were identified than in the previous Moran’s I test (Figure 4). Strong autocorrelation in the Peak District/Manchester region was detected in both tests. In the Hampshire/Hampshire region, a contrasting pattern emerged. This region showed strong agreement between Chi-square similarity and WHPT ASPT O/E, with both worsening over time. However, Hellinger similarity was increasing in regions with worsening WHPT ASPT O/E. Similarly, the Midlands region shows a region of disagreement between Hellinger similarity and WHPT ASPT O/E, with WHPT ASPT improving whilst Hellinger similarity worsens.

**Figure 4:**
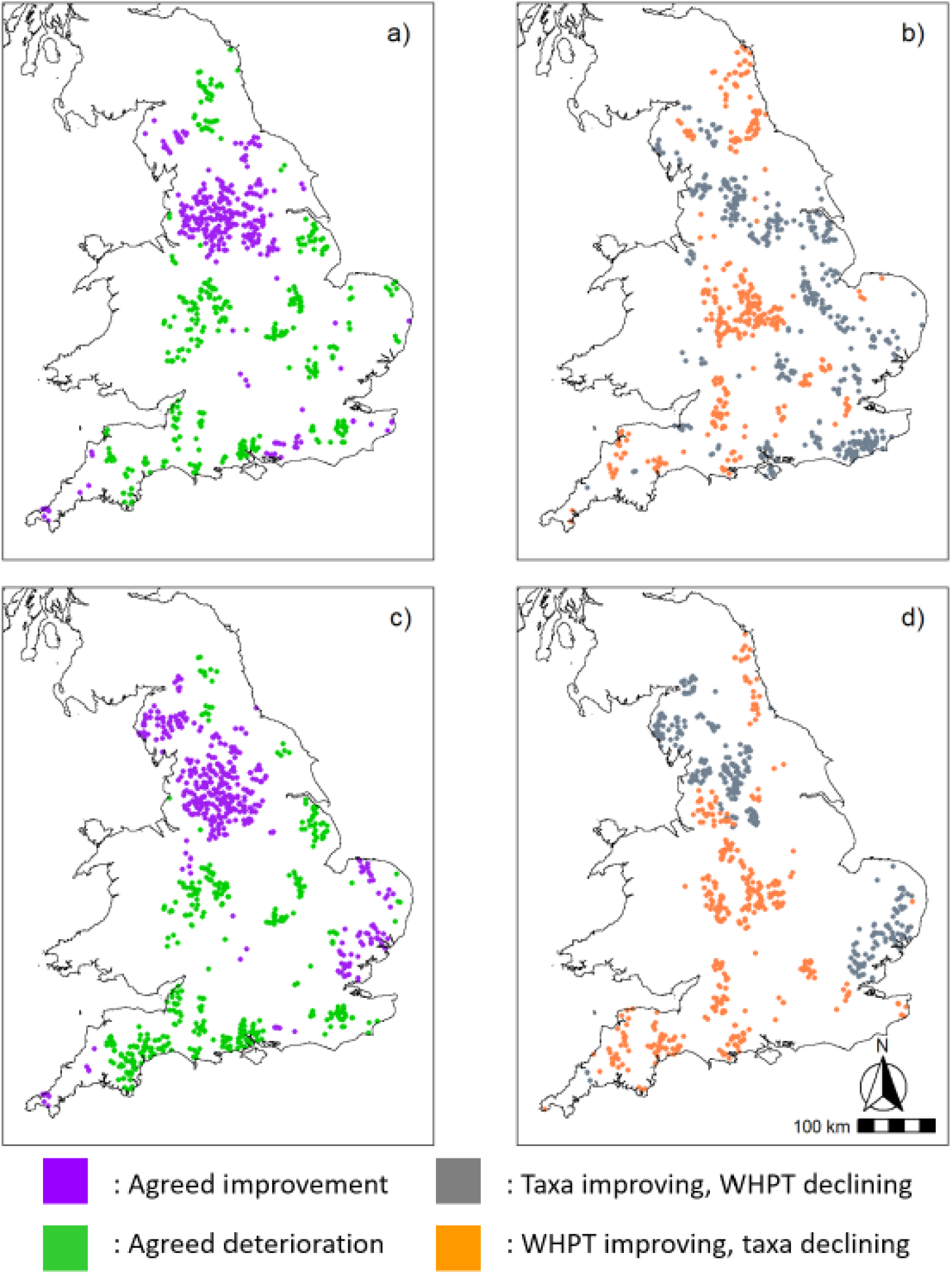
Spatial autocorrelation in regions of agreement and disagreement between WHPT ASPT and the similarity metrics. Maps a and b show agreement and disagreement between WHPT ASPT and Hellinger similarity respectively, and c and d show agreement and disagreement between WHPT ASPT and Chi-Square similarity. In maps a and c, green represents improvement and purple represents worsening. In b and d, grey represents the similarity metric improving whilst WHPT ASPT worsens, and orange represents the opposite

To test whether these patterns represented detectable changes in invertebrate assemblage composition, three key regions were identified and investigated more closely (Figure 1). These regions were selected due to the number of sites within each region, and for the Hampshire region due to the close clustering of sites. Regions such as the Essex/Suffolk area showed strong autocorrelation, but this was between more spatially dispersed sites. A PCA of the composition at each region over the past 20 years identified clear trends in composition underpinning the shifts in taxonomic similarity and WHPT ASPT O/E (Figure 5). The Lancashire PCA was characterised by a consistent shift along PC1 over a 20-year period. This was driven by increases in Elmidae, Hydropsychidae, and Gammaridae, and decreases in Chironomidae, Baetidae, and Asellidae. The West Midlands region expressed a gradual increase on PC1 between 2000 and 2009, before this trend reversed and the region underwent a marked negative shift on PC2. The initial trend was characterised by increases in Gammaridae, and declines in Caenidae, Hydrobiidae, and Chironomidae before partially reversing. The later shift was driven by increases in Sphaeriidae, Elmidae and Hydropsychidae, as well as a continuing decrease in Chironomidae. Average WHPT ASPT O/E ratios in this region were comparable to the Lancashire region. In the Hampshire region, there was minimal variation in the average composition of sites between 2000-2014. Between 2015 and 2019 there was a rapid shift away along the first and second principal components. This was characterised primarily by declines in Gammaridae, Hydrobiidae, Asellidae and Chironomidae, and increases in Elmidae, Oligochaeta, Baetidae and Sphaeriidae.

**Figure 5:**
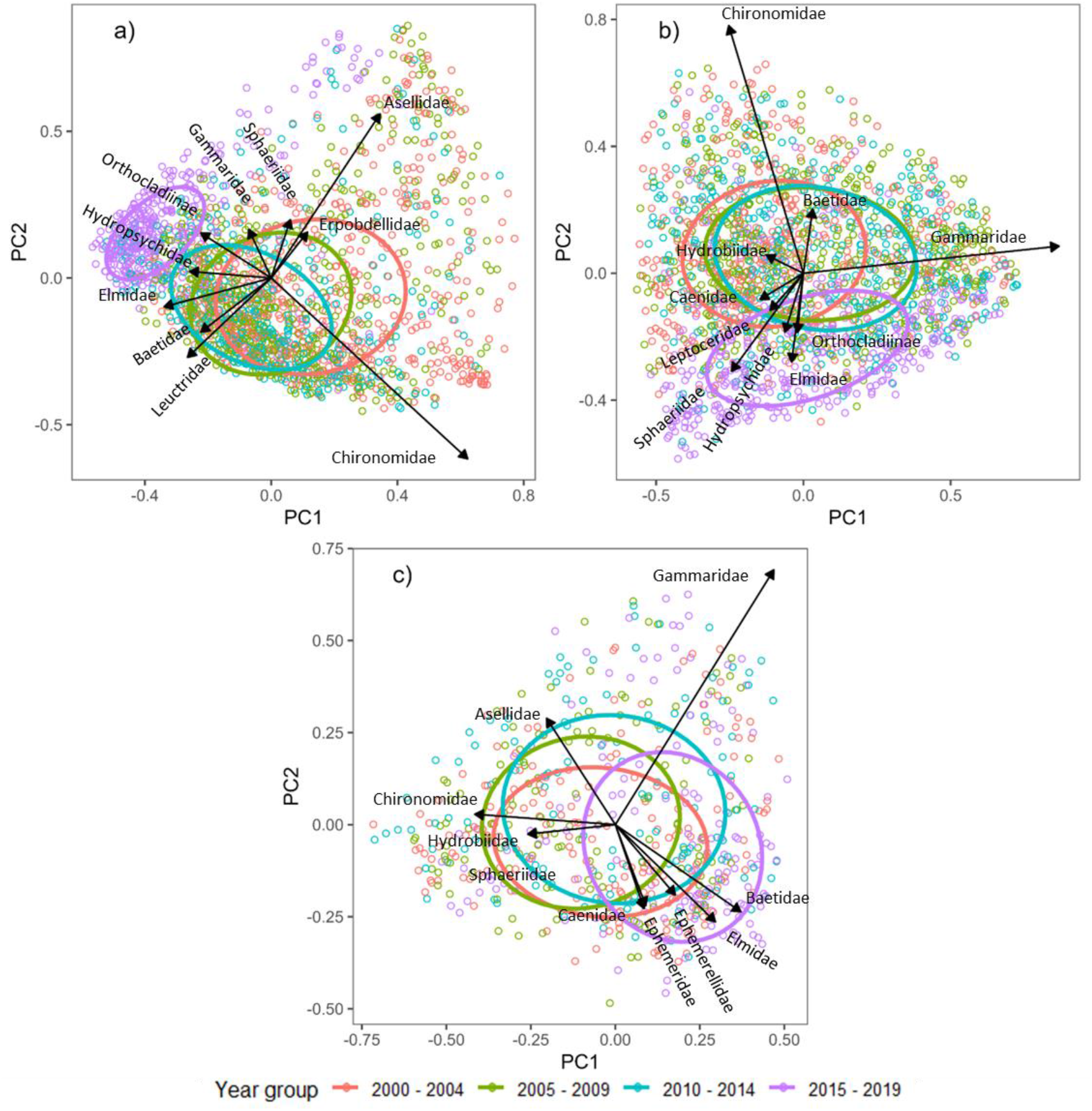
Principal Component Analysis undertaken for (a) Lancashire; (b) West Midlands; and (c) Hampshire regions (2000-2019), using 5-year aggregates to visualise change over time.

## Discussion

This study provides a national assessment of English river taxonomic trends relative to predictions derived from RIVPACS type models. WHPT ASPT O/E ratios were found to be improving overall at the national scale, with approximately 63% of all tested sites showing positive or marginally positive trends over time. In contrast, it was found that the majority of sites were moving away from their RIVPACS taxonomic predictions. When considering Chi-Squared and Hellinger similarity, 50.2% and 57.3% of sites showed trends away from their predicted assemblages, respectively. Notably, the integration of taxonomic trends and spatial analyses was able to detect regions with significant underlying shifts in composition that were undetected by WHPT ASPT O/E alone. Accordingly, the use of RIVPACS taxonomic predictions potentially offers a powerful and interpretable new tool for the management of freshwater environments.

### Are the target compositions achievable in a changing environment?

Most sites displayed trends away from RIVPACS taxonomic predictions over 10-years, in contrast to the most common metric (WHPT ASPT) used to assess ecological quality. There are several factors that could cause this pattern, either through limitations in the underlying RIVPACS model, or through the different patterns of recovery from multi-dimensional taxonomic data. For example, it is likely that taxonomic data are sensitive to multiple stressors simultaneously (Sarkis *et al*., 2023). Alternatively, the pathway to recovery that a taxonomic community travels might not result in linearly increasing similarity scores (Matthews *et al*., 2013; Matthews and Marsh‐Matthews, 2016). Intermediate compositions might be equal distance from the ideal communities as more impacted communities. Further work is required to assess the similarity scores of ecosystems in different ecological quality bands in a single year, to better understand the pathway to recovery for community compositions.

It is also possible that ramp stressors such as climate change limit the applicability of the reference site methodology used by RIVPACS-type models. As key environmental conditions such as precipitation and temperature continue to change, this will have direct impacts on the habitat suitability of sites for individual taxa (Vaughan and Gotelli, 2019). Indeed, it would be expected that the compositions at reference sites will change in line with antecedent environmental conditions. If the underlying RIVPACS model is not updated, this could result in predicted compositions that are unlikely to develop. This could be mitigated by regular updating of the RIVPACS reference site database to ensure regular resampling of reference sites and rerunning the underlying models using historical and current samples. Alternatively, an alternate framework could be developed by identifying key taxa in each defined RIVPACS ‘End-Group’ and running stacked species distribution models for each taxa based on habitat preferences from each taxon’s full range (Miller, 2010; Pollock *et al*., 2014). Additionally, models could be created that directly incorporate anthropogenic stressors (Wilkes *et al*., 2024). This would allow greater confidence in predicting the suitability of each new site for species and be more robust in accounting for the impact of climate change on abundances. Furthermore, the current tool could be used to assess the predictive performance in reference sites over time. As the latest update of the RIVPACS reference sites was in 2002, predictive performance over time with updated samples could be assessed to determine whether the impact of anthropogenic climate change or cyclical climatic phenomena (Milner *et al*., 2006) are altering baseline reference conditions.

Whilst most sites showed trends away from their taxonomic predictions, this was more pronounced for Hellinger similarity compared to Chi-squared similarity. The observed trend contrasts with recently observed trends between occupancy of rare and common taxa for UK species. Outhwaite et al. (2020) observed that in all tested clades over a 40-year period, rare and common taxa have had comparable increases in occupancy rate. However, occupancy was measured as change relative to 1970. Accordingly, changes in occupancy rates might be masked by longer term shifts that impacted a species range before 1970. Work undertaken by Tonkin et al. (2014) found that the distance between sites, and the overall occupancy of a regional taxa pool, were the largest determinants of colonisation by taxa. Accordingly, rarer taxa, that are often more sensitive to habitat degradation (Paisley *et al*., 2014), might have lower overall occupancy rates. If rarer sensitive taxa have been largely extirpated or reduced to low levels this could limit the recovery rate of sites that are undergoing improvement (Olden *et al*., 2004). Rather than measuring overall occupancy, this study has assessed the long-term trends of rare and common taxa towards RIVPACS predictions. The observed trends could be generated by RIVPACS having a lower predictive accuracy for rarer taxa through false negative occupancy from samples in the reference dataset and novel samples (Zhang *et al*., 2020). This would result in inflated similarities between observed and expected communities as the real trends of rare taxa are under sampled. This inflation could be mitigated by modifying the original RIVPACS model to a species-specific occupancy-based model, rather than a community-based model (Bailey *et al*., 2014). Further work is required to investigate the trends in rarer taxa recovery in English rivers, and to determine RIVPACS predictive accuracy for rarer taxa. Additionally, when using RIVPACS as a taxonomic target for management purposes, it is necessary to consider what metrics of ecological similarity, such as rarer or common taxa health, are the best indicators of overall ecosystem health. As prior studies have demonstrated that predicting rarer taxa is more challenging using RIVPACS-type models, it is likely that Hellinger similarity will be a more robust indicator of overall trends.

The Lancashire region showed major improvements in both WHPT ASPT O/E and taxonomic metrics between 2010-2019. This was driven by a gradual increase in the abundances of Elmidae, and Hydropsychidae, and decreases in Chironomidae and Baetidae. These changes are indicative of a region improving from a low to a medium ecological quality. Recent work has highlighted widespread improvements in macroinvertebrate communities driven by improvements in water quality up to 2015 (Pharaoh *et al*., 2023; Vaughan and Gotelli, 2019), which are more pronounced in urban streams than in rural streams. The agreement between pollution metrics and taxonomic metrics indicates that RIVPACS taxonomic predictions could infer underlying family-level trends as well as gaining traditional metric-level insights, thus providing greater potential for understanding the ecological processes driving recover. A key advantage of the RIVPACS methodology is the ability to generate targets and O/E ratios, allowing comparisons between highly varied sites to prioritise management strategies (Clarke *et al*., 2003; Wright *et al*., 1998). A current limitation in methods assessing taxonomic stability or improvements is this lack of comparability. Whilst urban streams might show greater rates of improvement, this could be a product of being in a lower condition initially. The current methodology partially mitigates this by allowing comparisons to simulated communities. This could be expanded to allow predictions and comparisons of species or family richness, functional diversity, or other metrics of taxonomic health, allowing greater comparison between rivers to better direct management efforts and identify key stressors.

### Sensitivity of taxonomic metrics to multi-stressors and novel stressors

A greater proportion of sites showed improvement in WHPT ASPT O/E compared to taxonomic similarity measures. Additionally, several regions were identified with contrasting trends in WHPT ASPT O/E and taxonomic similarity. WHPT ASPT O/E measures overall pollution sensitivity at a site, as well as fluctuations in oxygen availability (Paisley *et al*., 2014). Species that are less sensitive to pollution, such as those that can live in eutrophic waters, have lower scores than more sensitive taxa. There is functional overlap both between individual monitoring metrics, as well as between the scores each individual taxa represents (Jones *et al*., 2023). Improvements in WHPT ASPT O/E can be seen whilst a site might be impacted by stressors affecting LIFE O/E or the proportion of sediment-sensitive invertebrates (PSI O/E), measuring flow and sediment associations, respectively. Accordingly, this trend could be driven by sites that are significantly limited by multiple stressors, impacting the health of the macroinvertebrate community (Romero *et al*., 2018; Marshall and Negus, 2019). As pollution levels improve in these regions, other stressors could become more pronounced, or limit overall recovery. Management metrics often autocorrelate, for example reduced flow can impact oxygen availability, which in turn decreases WHPT O/E ratios. The use of multiple metrics simultaneously, in combination with taxonomic based metrics, could provide a clearer understanding of the key stressors at individual sites. Thus, the use of a single taxonomic metric of health sensitive to multiple stressors simultaneously is required (Feld and Hering, 2007). Additionally, taxonomic metrics of health will be sensitive to stressors to which metrics do not exist, such as the impacts of invasive species or pulse disturbance events such as floods or droughts (Guareschi *et al*., 2021).

Differing trends in WHPT ASPT O/E and taxonomic metrics were detected in multiple regions, with a large number of such sites detected in the Midlands. In this region, WHPT ASPT O/E was found to be improving whilst Hellinger similarity was found to be worsening overall. This was largely driven by decreases in Chironomidae, with increases in Sphaeriidae, Elmidae, and Hydropsychidae. This trend could be indicative of a region experiencing multi-stressor interactions, whilst one stressor has alleviated, driving changes in the underlying composition towards other impacted communities. Previous studies have highlighted that regions experiencing recovery in individual stressors will not recover to their maximum potential in the presence of alternate stressors, such as flow conditions or modifications (García-Barreras *et al*., 2023; Waite *et al*., 2020). When considering taxonomic similarity to RIVPACS predictions, this could result in improvements in individual macroinvertebrate metrics due to functional redundancy in the expression of metric scores, whilst showing no trend or decreases in taxonomic metrics. Whilst traditional metrics are largely one dimensional, with scores either improving or worsening, taxonomic metrics based on community composition are multi-dimensional (Matthews *et al*., 2013). Ecosystems can shift through a range of different compositions, and can establish multiple intermediate states when recovering or in the presence of alternate stressors (Chang and Turner, 2019). Whilst RIVPACS can generate a broad range of predictions for individual sites, these predictions are for rivers that have the lowest anthropogenic stressors for their type at the time of sampling. Accordingly, intermediate states in improving sites might show no overall improvement in taxonomic compositions. Additionally, as most rivers in England are impacted by anthropogenic stressors, it is possible that some sites might never achieve their targeted compositions (Vaughan and Ormerod, 2012; Kelly *et al*., 2008). Future work might consider comparing static scores in taxonomic similarity to multiple traditional metrics, to determine what pathways to recovery for taxonomic similarity are. Additionally, future iterations of RIVPACS-type models might consider being derived from all available data and incorporating information on key stressors, to be better able to predict the maximum potential for sites that experience unimprovable stressors such as the presence of urbanisation.

Contrasting taxonomic trends were detected in regions of England between Hellinger and Chi-Squared similarity. This was prevalent in the Hampshire region, which demonstrated increases in Hellinger similarity, but decreases in WHPT ASPT O/E and Chi-Squared similarity. This shift was driven by declines in Gammaridae, Hydrobiidae, Asellidae and Chironomidae, and increases in Elmidae, Oligochaeta, Baetidae and Sphaeriidae. This region comprises a range of semi-natural and urban streams, as well as a high proportion of chalk streams. The observed decline in Gammaridae has previously been detected across the south of England and Europe (Montgomery *et al*., 2022). Whilst it is likely that there are multiple stressors driving this shift, a prime candidate for this decline has been competition and extirpation through competition with invasive Gammaridae taxa (Cancellario *et al*., 2023). This region was one of the first sites that the species *Gammarus fossarum* was detected in Great Britain (Blackman *et al*., 2017). As the invasive Gammaridae taxa is not included in the RIVPACS predicted sites composition, this would result in decreases in similarity as expected Gammaridae were replaced with unexpected species. This suggests that RIVPACS taxonomic predictions might be capable of detecting the impact of novel stressors and contextualising these as impacts on the overall health or quality of an ecosystem. This is similar to previous studies which demonstrated that taxonomic metrics are as capable as traditional metrics in detecting the impact of invasive species (Guareschi *et al*., 2021). However, care must be taken to consider what is considered a healthy ecosystem. Hellinger similarity expected that sites in this region were improving in health overall as the abundances of more common taxa became more similar to expected communities, whilst Chi-squared similarity adequately detected that the abundances of rarer taxa were being impacted. However, being able to compare to static predicted baselines will allow for a clearer understanding of the impact of invasive taxa. The use of multiple baseline metrics will allow practitioners to rapidly detect novel stressors.

### Limitations and future work

This study investigated trends in RIVPACS derived taxonomic metrics and WHPT ASPT O/E over a 10-year period at sites in England, finding that a minority of sites are moving towards predicted targets for both taxonomic similarities, whilst most sites are improving in terms of WHPT ASPT O/E. The use of site-specific trends provides several advantages, allowing a clear prediction of whether English rivers are improving or worsening over time, negating the impact of changing sampling distributions over time, and allowing spatial trends to be identified. However, this significantly reduces the number of sites that can be assessed, with sites that are undergoing routine monitoring being heavily favoured. Whilst this was partially negated by lowering the criteria to 5 samples over a 10-year period, known biases exists in the dataset, such as many sites being routinely sampled for drought monitoring, or targeted investigation at known stressor hotspots. Additionally, whilst investigating trends at sites provides an understanding as to whether the taxonomic predictions derived from RIVPACS-type models are achievable or realistic, more work is required to determine the path of different river types to their target compositions, such as the impact of multi-stressors, or intermediate improving states. Future work might consider using RIVPACS derived taxonomic targets produced from our Monte-Carlo simulation to undertake static assessments of all samples in a single year, increasing the total number of sites that can be assessed. This could be interrogated to determine how multi-stressors impact distance from predicted compositions, and how close sites can come to their target compositions. This could also be used to generate thresholds for high, medium, and low quality rivers as utilised in the Water Framework Directive and UK Environment Act (European Union, 2000; UK Parliament, 2021).

A consistent challenge with implementing taxonomic monitoring in the freshwater environment has been the naturally high rates of variability and turnover observed for each species, and separating the background statistical noise from the influence of stressors (Arce *et al*., 2014).

Our study finds two potential pathways to aid in analysis. First, comparing community structure directly to predicted structures from a simulated baseline can allow for comparisons between sites, and contextualising whether a site is improving or worsening in condition. Second, the spatial comparison of the key taxonomic and metric trends provides a greater toolbox of information for freshwater managers. By comparing the impact of stressors such as pollution or flow modification using traditional metrics, both spatially and temporally, with taxonomic metrics that are sensitive to unidentified stressors such as invasive species, managers would have a better understanding of the key limitations at sites and be able to monitor the development of novel stressors. This will allow for the development of more robust and reactive management strategies, or the quantification of management or restoration targets such as biodiversity net gain scaled to a maximum achievable biodiversity for each site (Gurnell *et al*., 2020; Wilkes *et al*., 2024).

The use of spatial autocorrelation analyses highlighted regions in which clear trends were present in the tested metrics. This allowed an exploration of key drivers and limitations in using RIVPACS derived taxonomic targets, such as identifying regions in which WHPT ASPT O/E and taxonomic targets are diverging. A key challenge with investigating large-scale datasets is identifying overall trends and spatial patterns in a meaningful manner so that drivers can be identified. This is particularly challenging in freshwater ecosystems, where naturally high rates of turnover and stochasticity increase background noise and variation (Le *et al*., 2021; Benke, 1998; Booth *et al*., 2016). However, many stressors that impact freshwater ecosystems can be spatially limited in scale and might only impact an individual river catchment or small reach, such as abstraction or point-source pollution. Thus, techniques relying on spatial autocorrelation will struggle to identify these stressors. It is recommended that a hybrid approach using larger scale spatial analysis which can detect trends due to land use, invasive species, or climate change, as well as catchment or reach scale trends, be deployed by managers.

## Conclusions

This study produced a novel Monte Carlo simulation to incorporate into RIVPACS-type models to allow for observations to be contrasted with taxonomic targets rapidly and efficiently. An assessment of 10-year taxonomic trends in England determined that only a minority of sites are moving towards their taxonomic targets as determined by RIVPACS predictions; however, this is more prevalent for common taxa compared to rare taxa. Additionally, strong agreement was identified between trends in WHPT ASPT O/E and taxonomic trends in some regions. This highlights that RIVPACS derived taxonomic targets can be a feasible management metric. Patterns of spatial auto-correlation were found in which WHPT ASPT O/E trends and taxonomic trends diverged. It is speculated that these are driven primarily by either non-linear paths of taxonomy towards their target compositions, or by the impact of multiple stressors limiting recovery at a site. Taxonomic targets were found to successfully detect widespread changes in composition driven by changes in underlying conditions, and might be sensitive to multiple uncharacterised stressors such as invasive taxa. Accordingly, the use of taxonomic targets can expand the toolbox available to freshwater managers, allowing novel or worsening stressors, such as climate change, invasive species, or pollutants to be readily identified. It is recommended that RIVPACS derived taxonomic targets be considered as a new measure of assessing river ecosystem health, and enabling new approaches to understanding what pathways to ‘good’ or ‘excellent’ status should look like in rivers with multiple stressors.

## Acknowledgments

This work was funded as part of a Natural Environment Research Council (NERC) CASE Studentship (NE/R007853/1). Lee E. Brown also acknowledges support from the NERC Drivers and Repercussions of UK Insect Decline (DRUID) project (NE/V006916/1)

